# MetaLATTE: Metal Binding Prediction via Multi-Task Learning on Protein Language Model Latents

**DOI:** 10.1101/2024.06.26.600843

**Authors:** Yinuo Zhang, Phil He, Ashley Hsu, Pranam Chatterjee

## Abstract

The bioremediation of environments contaminated with heavy metals is an important challenge in environmental biotechnology, which may benefit from the identification of proteins that bind and neutralize these metals. Here, we introduce a novel predictive algorithm that conducts **Metal** binding prediction via **LA**nguage model la**T**en**T E**mbeddings using a multi-task learning approach to accurately classify the metal-binding properties of input protein sequences. Our **MetaLATTE** model utilizes the state-of-the-art ESM-2 protein language model (pLM) embeddings and a position-sensitive attention mechanism to predict the likelihood of binding to specific metals, such as zinc, lead, and mercury. Importantly, our approach addresses the challenges posed by proteins from understudied organisms, which are often absent in traditional metal-binding databases, without the requirement of an input structure. By providing a probability distribution over potential binding metals, our classifier elucidates specific interactions of proteins with diverse metal ions. We envision that MetaLATTE will serve as a powerful tool for rapidly screening and identifying new metal-binding proteins, from metagenomic discovery or *de novo* design efforts, which can later be employed in targeted bioremediation campaigns.

## Introduction

Metals play a crucial role in natural systems, serving as co-factors for the functionality of enzymes and catalysts for chemical reactions and pharmaceutical production [Kostenkova et al., 2022]. Transition metals are essential due to their wide range of oxidation states and involvement in various biological processes. Zinc is a key component of zinc finger proteins, which are involved in DNA binding and transcriptional regulation [Klug, 2010]. Most of the proteins in the Protein Data Bank are metalloproteins [Permyakov, 2021], underscoring the ubiquity of metal-protein interactions in biology. However, certain heavy metals, such as cadmium, lead, and mercury, can be toxic to living organisms at high concentrations [Tchounwou et al., 2012, Chen et al., 2023b]. These toxic non-essential heavy metals can disrupt cellular functions by interfering with essential metal-dependent processes. For example, they often exhibit higher affinity for sulfhydryl groups in functional metalloenzymes and sometimes possess structurally-similar functional groups compared with essential metals like zinc, copper, and iron [Tripathi and Poluri, 2021]. Such metal replacements can adversely affect cells by disturbing membrane transport proteins, reducing enzymatic activities, and promoting the accumulation of reactive oxygen species (ROS), ultimately leading to disrupted metabolic activities [Tripathi and Poluri, 2021].

The ability to predict metal-binding properties of proteins is crucial for several reasons. First, it may enable the engineering of remediating proteins to sequester and detoxify heavy metals from contaminated environments [Dixit et al., 2015]. This is particularly important given the persistent and non-biodegradable nature of heavy metals, which necessitates their continuous removal from soil and water bodies to maintain ecological balance [Permyakov, 2021, Chen et al., 2023b]. Additionally, with the increasing availability of genomic data from understudied organisms, such as mollusks, predicting metal-binding properties of novel proteins can provide insights into their roles in these organisms and their potential for biotechnological applications [Calatayud et al., 2021].

Existing metal predictor tools, such as MIB2 [Lu et al., 2022], require the user to provide structural information in PDB format to predict binding sites. However, many proteins from understudied organisms either do not have canonical structures solved or are structurally disordered, precluding usage of structure prediction methods such as AlphaFold [Abramson et al., 2024, Calatayud et al., 2021, Permyakov, 2021]. Even for many of the current known metal-binding proteins, their status *in vivo* is still unknown [Permyakov, 2021]. Recently, protein language models (pLMs) like ESM-2 and ProtT5 have shown the ability to capture evolutionary and functional information from sequence information alone, without the need for structural data [Lin et al., 2023, Elnaggar et al., 2021]. Previous studies have utilized pLM embeddings as input for classification tasks, such as predicting protein cellular localization and signaling [Thumuluri et al., 2022, Teufel et al., 2022]. Other works have utilized pooled protein embeddings for ion binding site predictions, but are largely restricted to common ions [Shenoy et al., 2024, Yuan et al., 2022] and overlook the imbalanced nature of metals in current datasets. These models further tend to perform separate predictions per metal, which does not align with the nature of metal-binding proteins, as they usually bind to two or more ions simultaneously [Permyakov, 2021], or interchangeably with pH changes [Bofill et al., 2009].

To this end, we present a **Metal** binding predictor using protein **LA**nguage model la**T**en**T E**mbeddings from ESM-2. MetaLATTE can input any protein sequence and predict the likelihood of binding to an array of diverse metals, including those that are underrepresented and considered environmentally toxic. Via a multi-task learning strategy with elements borrowed from image generation [Schroff et al., 2015], MetaLATTE provides a probability distribution over a diverse set of metals for an input protein sequence. Our results demonstrate that MetaLATTE can accurately identify likely metals with which a protein interacts, from sequence alone. Overall, our work highlights the immense utility of pLMs for protein function prediction, and motivates downstream metal-binding protein design efforts.

## Methods

### Dataset Preparation

#### A. Stage 1 Data Preparation

Training data for metal-binding proteins was obtained from the MbPA database [Li et al., 2023]. To focus on transition and heavy metals, we selected proteins binding to silver (Ag), cadmium (Cd), cobalt (Co), copper (Cu), iron (Fe), mercury (Hg), Manganese (Mn), molybdenum (Mo), nickel (Ni), lead (Pb), platinum (Pt), vanadium (V), tungsten (W), and zinc (Zn) (Figure S1). Metal labels with fewer than 6 samples were excluded to ensure sufficient representation for each metal class. Obvious negative samples, i.e., proteins not binding to any metal, were obtained from the Mpbipred database [Aptekmann et al., 2022] to provide a non-binding label, and thus form a balanced training set for the multi-label classification task.

The dataset was split into training and validation sets using balanced stratification separation [Sechidis et al., 2011] to ensure a fair distribution of metal labels across the folds. This approach was employed for 4-fold cross-validation training to assess the model’s performance and generalization ability. The test dataset consisted of two parts: (1) a non-overlapping dataset from MetalPDB [Andreini et al., 2012], which was curated to include metal-binding proteins not present in the training data, and (2) newly sequenced mollusk genomic sequencing results [Calatayud et al., 2021] that were not documented in any protein database at the time of the study. As a note, it is worth mentioning that as some of the transition metals are heavily understudied (Pb, V, for example), the held-out test set does not contain protein sequences that bind to those metals and is dominated by those that interact with well-studied metal ions like Zn and Mn.

#### B. Stage 2 Data Preparation

For Stage 2 training, we focused the model to learn site-specific information. To do this, we created decoy protein sequences by modifying the binding site amino acids in the original metal-binding proteins. The most distant amino acid to the binding amino acid, according to the BLOSUM62 matrix [Henikoff and Henikoff, 1992], was used to replace the original amino acid spanning the binding site with a *±* 3 residue window. This resulted in the original metal-binding proteins containing contaminated amino acids flanking the binding motif sites. These decoy sequences serve as “semi-hard” negative samples for the model. We also created a “hard” negative dataset in which we performed alanine swapping to the binding sites labeled by MbPA database.

As training data for the second stage of the model, we created triplet batches using the original metal-binding proteins, their corresponding decoy negative sequences, and additional positive sequences. For each anchor sequence, we identified a positive sequence that shared metal-binding labels, along with its corresponding decoy sequence, forming an [anchor, positive, negative] triplet for triplet loss minimization. The triplets were pre-batched dynamically based on sequence lengths to optimize memory usage and training efficiency. After certain epochs, we gradually swap the semi-hard negatives with hard negative datasets to further enable the model to distinguish between nuances within the embeddings that related to possible function loss.

To address class imbalance, we applied an oversampling strategy for minority labels. Specifically, for anchor sequences with rare metal-binding labels (e.g., Cd, V, W), we generated multiple triplets by finding different positive sequences that share the same metal-binding labels. This oversampling approach increased the representation of minority classes within the training set, enabling the model to learn from a more balanced distribution of metal-binding labels. The dataset was then split into training and validation sets via a standard 0.8 to 0.2 ratio using stratified shuffle-split [Sechidis et al., 2011], ensuring that the distribution of metal-binding labels was preserved in both sets.

### Model Architecture

Since metals bind to specific charged amino acid motifs, we sought to train a model that is sensitive to positional information changes in proteins. We built our model on top of the pre-trained ESM-2-650M pLM [Lin et al., 2023], which has already acquired basic protein evolutionary information and is well-validated in the literature for both prediction and design tasks [Brixi et al., 2023, Bhat et al., 2023, Chen et al., 2023a, Su et al., 2023, Peng et al., 2024, Vincoff et al., 2024]. However, current pLMs tend to capture global evolutionary patterns instead of the site-specific motif changes required for domain-specific tasks. As such, we define the architecture in two stages:

#### A. Stage 1 Training

In the first stage of model training (Figure 1), we fine-tuned the model for multi-label classification tasks using the 14 heavy metal labels and a non-binding label. We unfroze the last two layers of ESM-2-650M to adjust its pre-trained embeddings. To capture site-specific information, we added an attention pooling layer and rotary position embeddings [Su et al., 2024] to transform the original embedding output before the classification head. The attention pooling layer helps the model focus on relevant regions of the protein sequence, while the rotary position embeddings provide positional information to the model.

**Figure 1:**
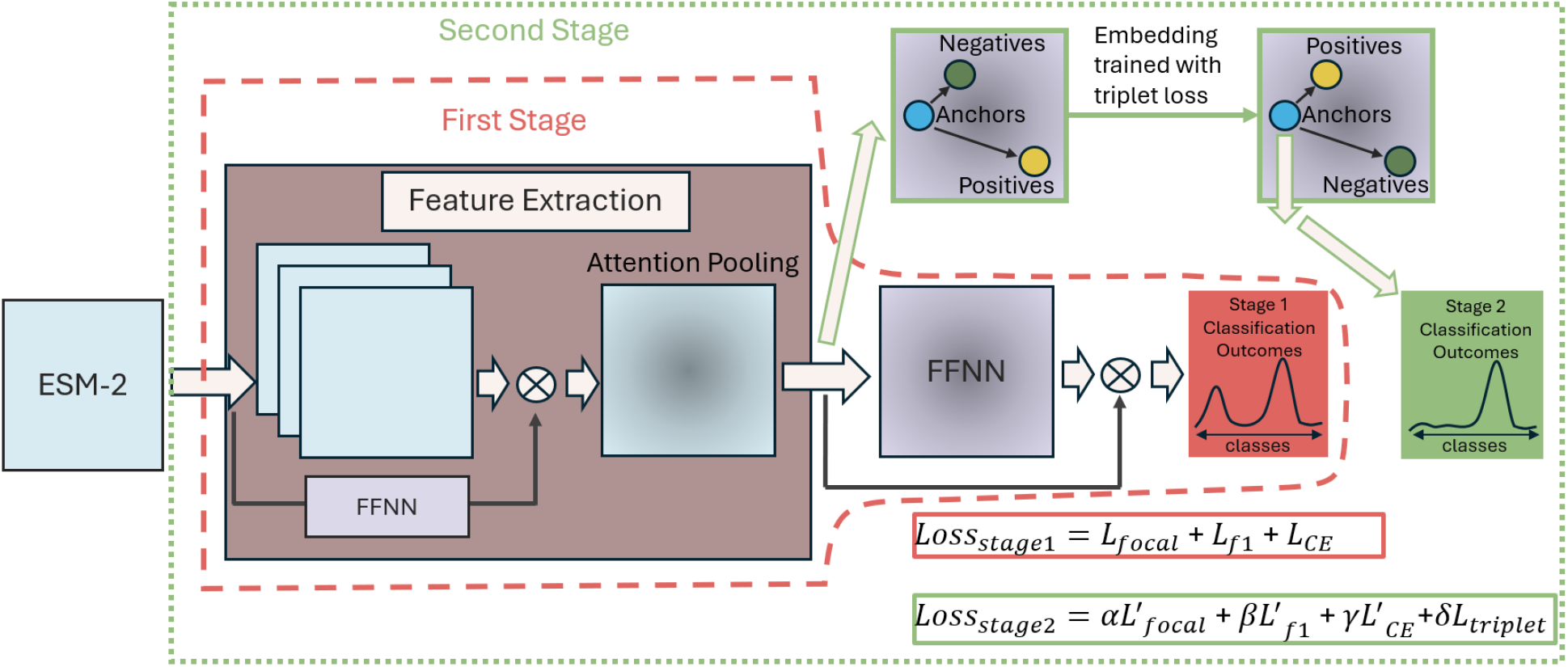
MetaLATTE schematic. (1) First Stage: An input protein’s high-dimensional ESM-2 representation is processed using attention pooling, followed by a classification head leveraging focal loss, F1 loss, and reconstruction loss for multi-label prediction. (2) Second Stage: MetaLATTE is further tuned with hard negatives and positives to form triplet batches, enhancing sensitivity to binding motifs via a combination of triplet loss, focal loss, F1 loss, and reconstruction loss.

We performed 4-fold cross-validation training and selected the best model based on the lowest validation loss. The classification thresholds were updated during training using exponential moving average (EMA) [Yan et al., 2022] with a historical memory factor (*α*) of 0.6, which determines the balance between the current and previous threshold values. The batch size was set to 6, and the peak learning rate was set to 1e-3 with a cosine learning rate scheduler with warm-ups of 200 steps [Touvron et al., 2023]. The entire training was completed within 32 GPU hours on an 7xNVIDIA A6000 DGX server with 350 GB of shared VRAM using the PyTorch Lightning framework for distributed data-parallel training [Falcon and The PyTorch Lightning team, 2019].

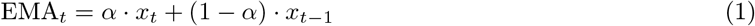

#### B. Stage 2 Training

For stage two, we continued training the model from stage one (Figure 1). We employed a cosine learning rate scheduler with a warm-up phase of 100 steps, reaching a peak learning rate of 2e-4 [Touvron et al., 2023]. The embeddings for the triplets were generated using shared model weights, enabling the model to learn a consistent representation space and capture the similarities and differences between the hard negatives and hard positives effectively.

To stabilize the model’s performance, we increased the historical memorization factor from 0.6 to 0.9, allowing the model to give more importance to the previously learned classification thresholds and gradually adapt to new information. This smoother update helps the model settle down with better classification thresholds. Stage 2 training was completed within 56 GPU hours on the 7xNVIDIA A6000 server with the same strategy as Stage 1 training except with a dynamically pre-batched dataset.

### Loss Construction

The contribution of each loss function to the model’s training is a tunable parameter. For Stage 1, we experimented with equal contribution among the focal, F1 loss, and reconstruction CE loss functions. For Stage 2, we attributed more weight to the triplet loss compared to the other two loss functions. The specific weights were determined through empirical experiments and validation performance.

#### A. Class Balanced Focal Loss

We used class balanced focal loss for both stages of training. Focal loss has shown its ability to train classifiers with minority classes [Cui et al., 2019]. The hyperparameter *γ* was set to 1.2 based on parameter tuning, and *β* was set to 0.9999 to adjust the class weights.

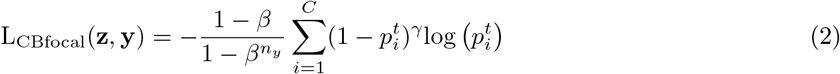

where

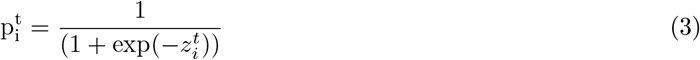

#### B. F1 Loss

We implemented a custom soft F1 loss function to optimize the model’s performance. This F1 loss calculates the true positives (TP), false positives (FP), and false negatives (FN) for each class based on the predicted probabilities and the ground truth labels. It then computes the precision and recall values for each class using these counts. By minimizing the F1 loss, we encourage the model to find a balance between precision and recall, leading to improved overall performance than focal loss alone.

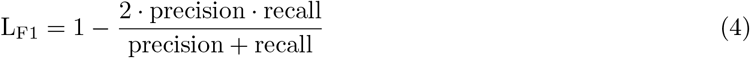

#### C. Reconstruction Loss

We also introduce reconstruction loss to the model for both stages to prevent the model from deviating from natural protein distributions. The reconstruction layers teach the model how to project embeddings back to logits with cross-entropy loss, enabling downstream sequence design in the future.

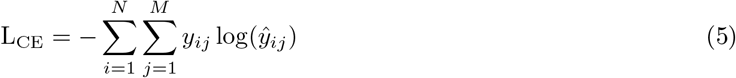

where *N* is the sequence length and *M* is the vocabulary size inherited from ESM-2 model.

#### C. Triplet Loss

Triplet loss, inspired by image classification tasks [Schroff et al., 2015, Balntas et al., 2016, Paszke et al., 2019, Liu et al., 2019], enables learning of nuanced differences in the embedding space that differentiate different labels. Considering that the negative samples we created were “hard” training examples, we adopted an adaptive margin in the loss equation to dynamically adjust the margin based on the distances between the anchor-positive and anchor-negative pairs. After epoch 10, semi-hard triplets are gradually replaced by hard triplets, to generate a better embedding representation for the protein sequences that are aware of mutations.

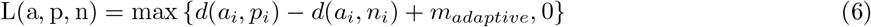

where *i* is the minibatch, *m* is the margin and

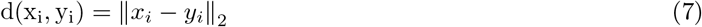

The adaptive margin *m*_*adaptive*_ is computed as:

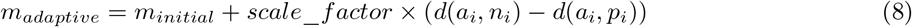

where *scale*_*factor* is set to 0.1. This approach ensures that harder triplets are assigned with larger margins, while easier triplets have smaller margins.

### Benchmarks

For multi-label classification results comparison, we used sensitivity, precision, recall, macro F1-score, Matthews Correlation Coefficient (MCC), Area Under the Receiver Operating Characteristic Curve (AUROC), and proportional accuracy score from module scikit-learn (v1.3.2) [Pedregosa et al., 2011] as evaluation metrics.

#### A. Embedding Source Comparison

We compared MetaLATTE’s performance with embeddings generated using VHSE [Mei et al., 2005] and one-hot encoding. An MLP classification head was trained on these embeddings using the same dataset as MetaLATTE, and the best model was selected based on validation loss.

#### B. Model Comparison

We benchmarked MetaLATTE against an XGBoost model, a state-of-the-art gradient boosting algorithm known for its strong performance on high-dimensional input data [Chen and Guestrin, 2016], optimized using optuna [Akiba et al., 2019] for hyperparameter tuning. The best XGBoost model, trained on ESM-2 default embeddings with EMA for threshold updates, was obtained after 200 trials of hyperparameter optimization. For mollusk sequences, sequence alignment was performed by LALIGN from EMBL-EBI [Pearson, 1991] and Geneious alignment, both via the BLOSUM62 matrix [Henikoff and Henikoff, 1992]. We used sensitivity to further evaluate classification model performances.

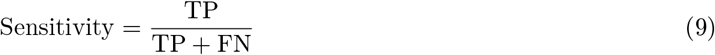

#### C. Visualization

We used UMAP with Louvain clustering from scanpy (v1.9.8) [Wolf et al., 2018] to visualize the shifting of learned embeddings space for the validation dataset. The number of neighbors parameter in UMAP was set to 50.

## Results

### MetaLATTE accurately predicts rare metal binding proteins

We first evaluated our model’s performance on available metal-binding proteins from MetalPDB [Andreini et al., 2012] that were not present in our training database MbPA [Li et al., 2023]. Preliminary results show that MetaLATTE, using ESM-2 embeddings, achieves higher scores for classification metrics compared to that of simpler embeddings, such as biophysical VHSE [Mei et al., 2005] vectors and one-hot encodings (Figure S2A). We hypothesize that the additional information regarding positions of amino acids in ESM-2 augments MetaLATTE’s ability to capture protein’s motif information critical for binding to rare metals.

To benchmark MetaLATTE, we compared it to XGBoost [Chen and Guestrin, 2016], optimized using optuna [Akiba et al., 2019]. Both models use ESM-2 embeddings and EMA [Yan et al., 2022] for threshold updates. While overall performance metrics were comparable (Figure S2B), MetaLATTE demonstrated superior capabilities in predicting rare metal binding. Figure S3 and Figure S4 illustrate the per-class performance on the validation set, revealing MetaLATTE’s consistent performance across all metals, including rare earth metals with small sample sizes. In contrast, XGBoost struggled with these rare classes (Figure 2). Table 1 provides a detailed comparison of sensitivity for each metal class:

**Table 1:**
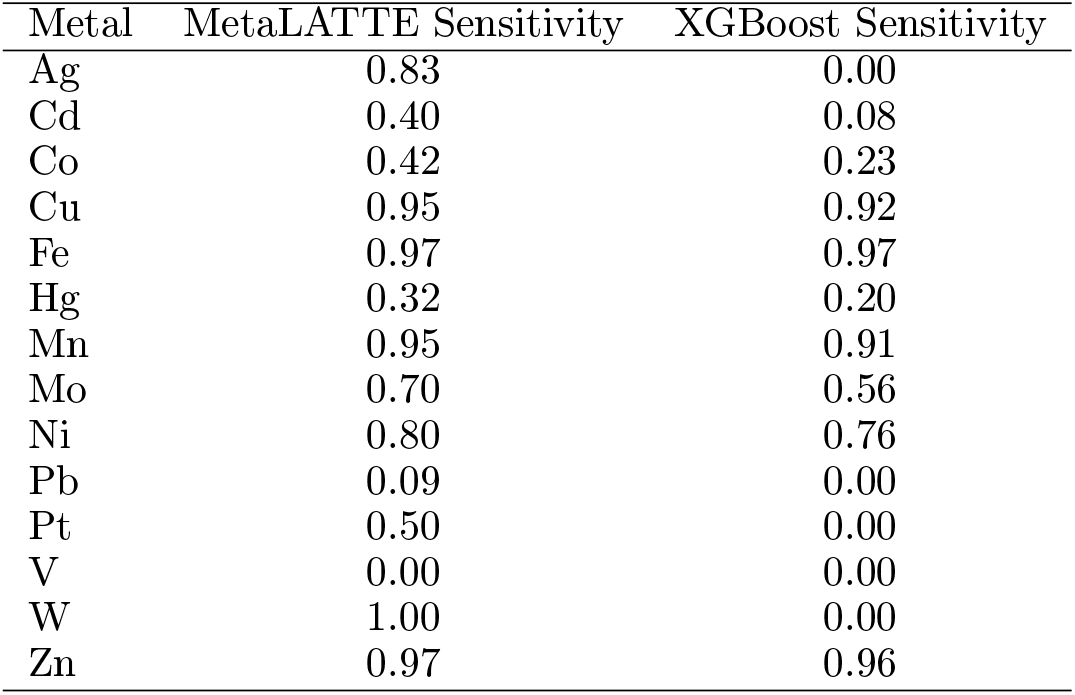
Sensitivity comparison for metal binding prediction.

**Figure 2:**
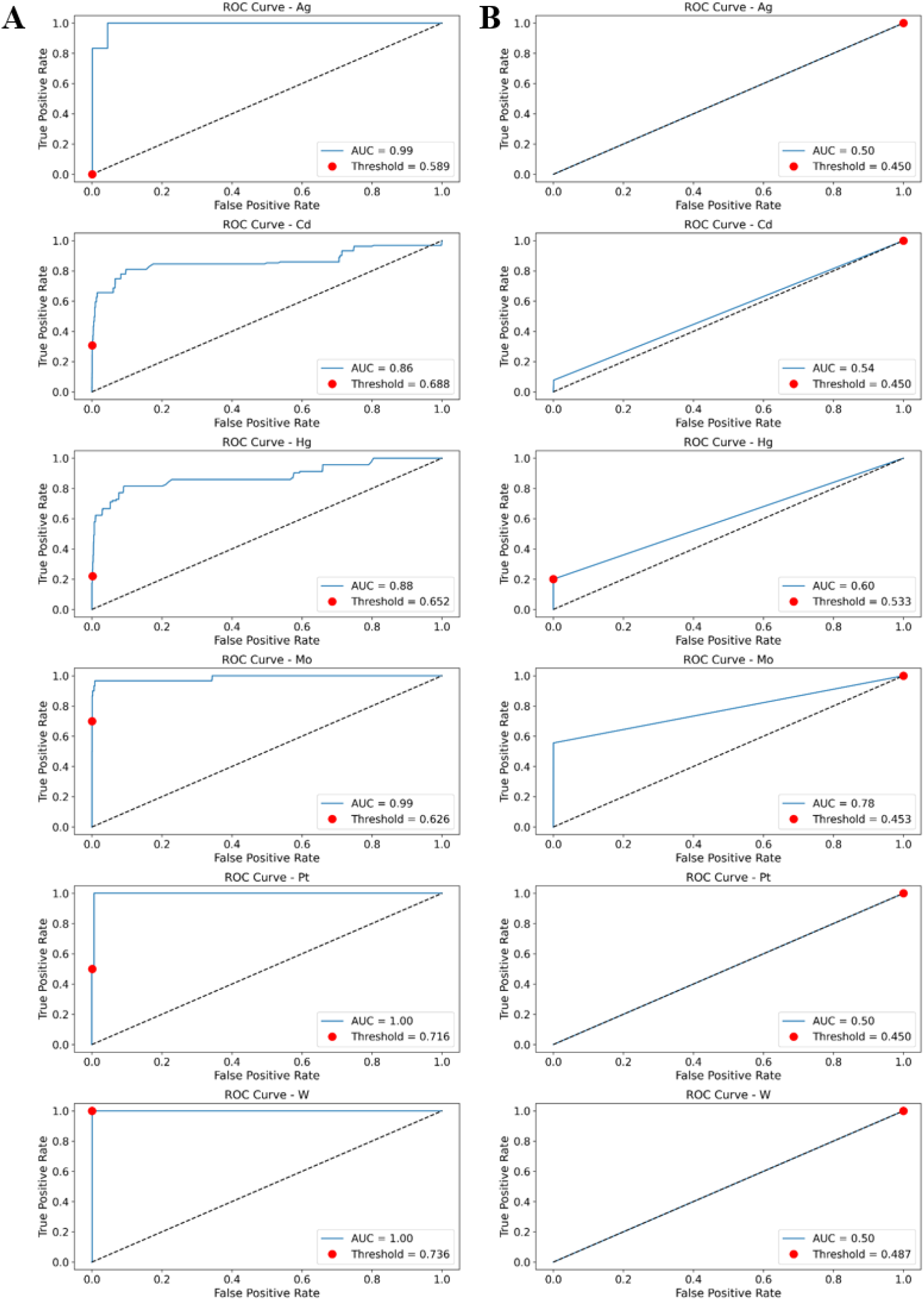
ROC curves for classification on individual metals. Classification thresholds are labeled as the red points for each metals depicted. Metals are indicated in the title of each plot for the (A) MetaLATTE Stage 2 model results, (B) XGBoost results.

Notably, MetaLATTE shows superior performance for rare metals, including Ag, Cd, Hg, Mo, Pt, and W. Our results highlight the challenge of training classifiers for rare metal classes and underscores MetaLATTE’s potential for environmental remediation applications that focus on binding these underrepresented metals. It is also worth noting that while overall test set metrics may appear similar between models, this is largely due to the dominance of common metals like Zn and Mn in the test set. MetaLATTE’s ability to maintain strong performance across almost all metal classes, including those that are underrepresented, demonstrates its robust and generalizable approach to metal binding prediction.

### MetaLATTE detects shift of embeddings from decoys

To further investigate MetaLATTE’s ability to capture subtle changes in metal-binding properties due to mutations in binding motifs, we visualized the embedding space of the triplet datasets using UMAP. As shown in Figure 3, UMAP visualization reveals a clear separation between the anchor proteins and their corresponding negative clusters, which represent proteins with mutated binding motifs, indicating that MetaLATTE identifies differences in the embedding space caused by these mutations, effectively predicting loss-of-function scenarios. While stage one focal loss alone can differentiate embeddings for different labels, the triplet training with introduced mutations around motif sites enables MetaLATTE to better distinguish functional changes due to amino acid mutation, suggesting improved adaptation to more generalized cases. Overall, the distinct clustering of anchor and negative proteins demonstrates that MetaLATTE’s learned embeddings capture nuanced changes in metal-binding properties. With further training on diverse proteins, we expect MetaLATTE to generalize to functional prediction for newly sequenced proteins, such as those from metagenomics studies. These results highlight MetaLATTE’s potential to both identify novel metal-binding proteins and model their functional roles in various biological processes.

**Figure 3:**
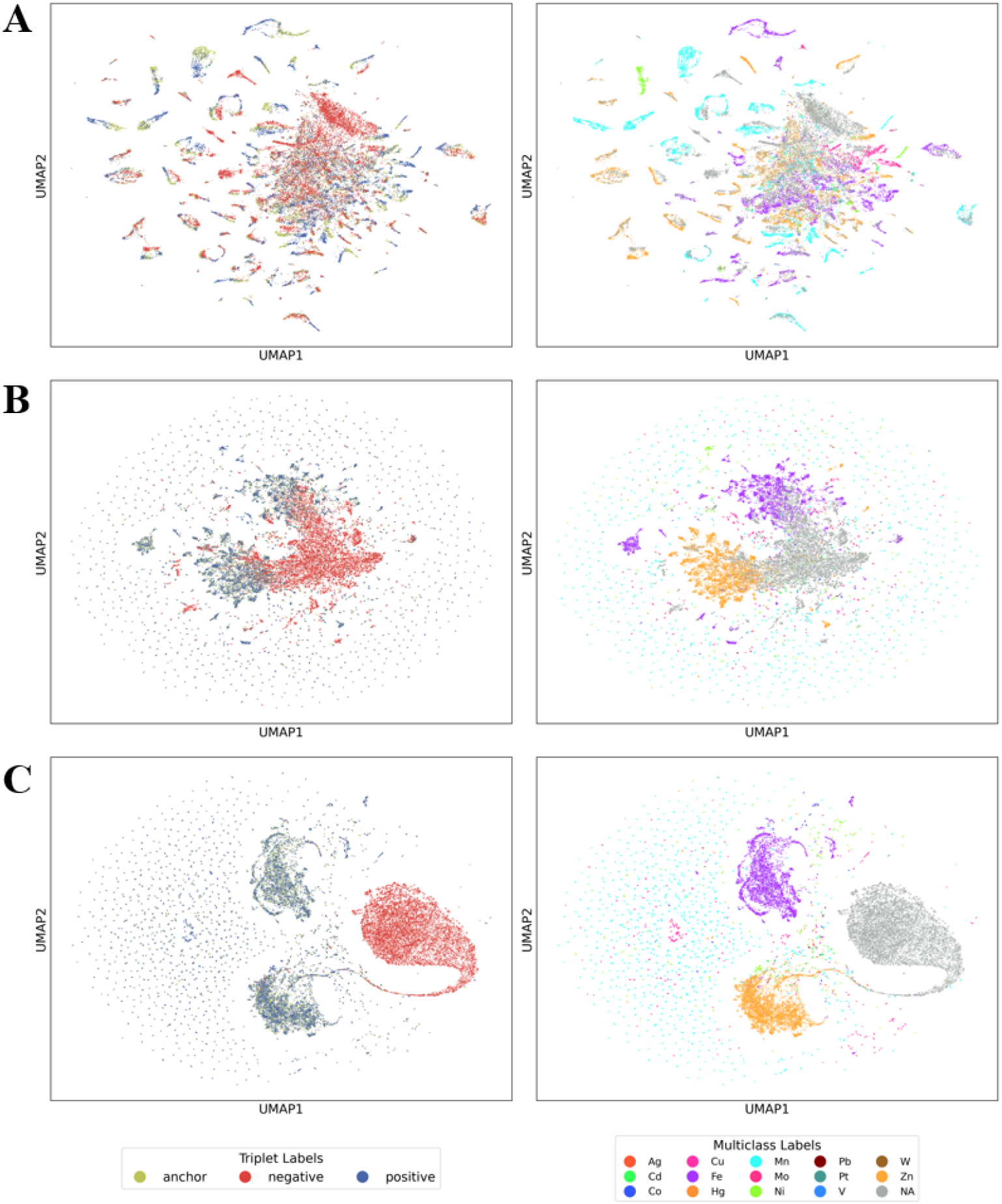
UMAP visualization of embeddings of the triplet dataset. UMAP plots on the left represent the embeddings colored by the anchor, negative, positive triplet labels, while plots on the right are the same embeddings colored by their actual metal binding labels. (A) ESM-2 Default Embedding for the Triplets (B) MetaLATTE Stage 1 Model Embedding for the Triplets (C) MetaLATTE Stage 2 Model Embedding for the Triplets.

### MetaLATTE predicts binding metals for understudied proteins

Finally, to assess MetaLATTE’s performance in a real-world scenario, we tested both stages of our model on proteins directly translated from the sequencing results of isoform RNAs in understudied mollusk *Helix pomatia* [Chabicovsky et al., 2003]. These short proteins play a crucial role in bioremediation by participating in heavy metal absorption in slugs and snails, which serve as sentinel organisms for measuring regional environmental pollution [Baroudi et al., 2020]. Despite their importance, these proteins are not present in any structural database. Rather, the protein expressions were validated via ion exposure experiments [Chabicovsky et al., 2003, Calatayud et al., 2021]. MetaLATTE successfully predicted Cu and Cd ion binding in specific poorly-structured isoforms, distinguishing among HpoMT1 (Cu-specific), HpoMT2 (Cd-specific), and HpoMT3 (binds both) (Figure 4A). Even though HpoMT1 and HpoMT3 share 75.4% sequence identity with pair-wise alignment, the nuanced amino acid changes and their corresponding functional shift is still picked up by MetaLATTE. Notably, MetaLATTE detected a loss-of-function (predicts “non-binding”) in the Cd-specific HpoMT2 protein when binding site amino acids were deliberately mutated (Figure 4B) to cysteines. These results demonstrate MetaLATTE’s superior ability to predict metal-binding properties of poorly-characterized proteins, highlighting its potential for discovering novel metal-binding proteins in understudied organisms.

**Figure 4:**
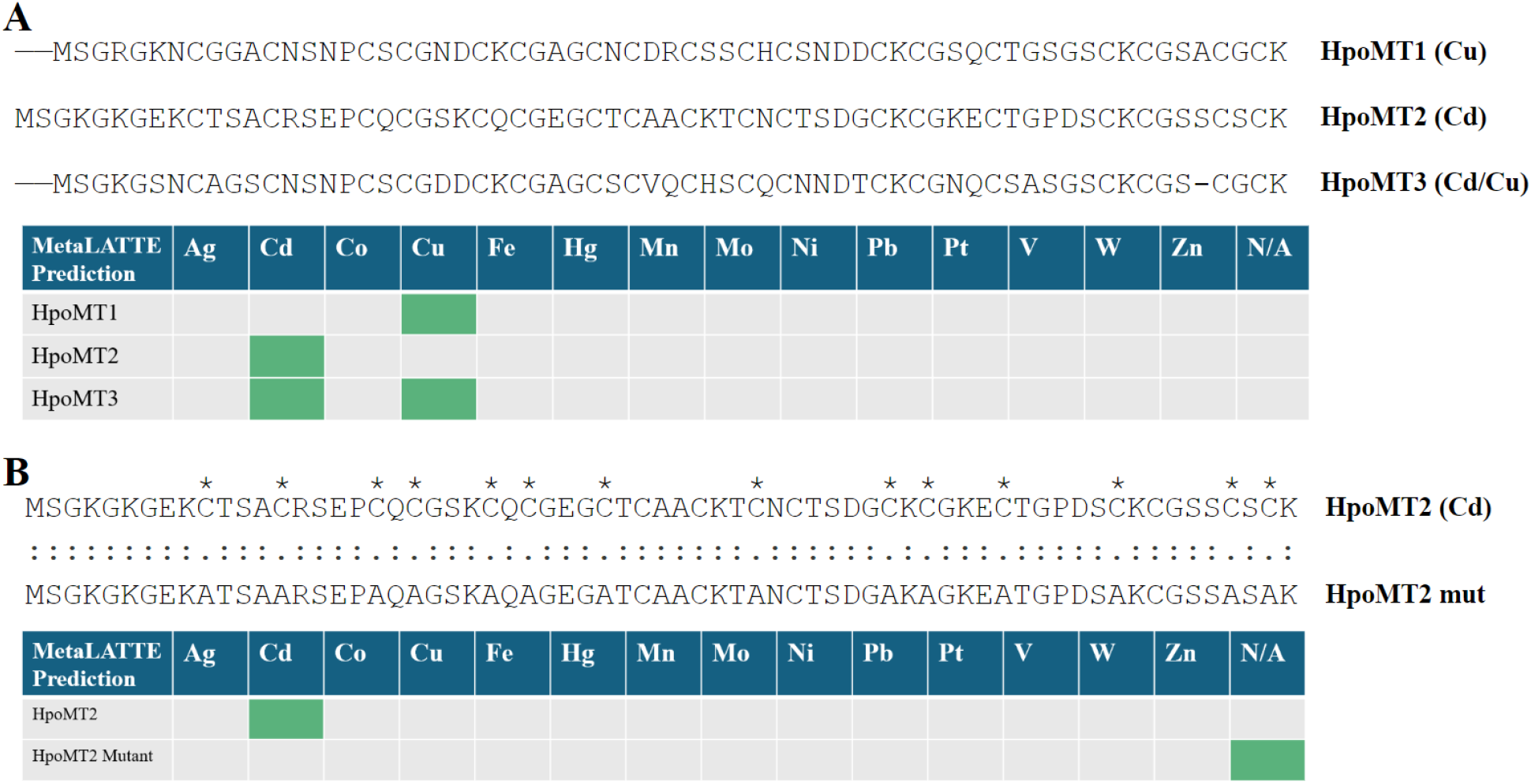
Multiple sequence alignment (MSA) and local pair-wise alignment results of HpoMT isoforms alongside MetaLATTE predictions. (A) HpoMT1, HpoMT2, and HpeMT3 are metallothionein isoforms in the snail *Helix pomatia*. Notably, HpoMT2 is a Cd-specific metallothionein, whereas HpoMT3 can bind to Cu ions as well. The table depicts which metals MetaLATTE predicts for interaction, provided the input protein sequence specified. (B) Loss-of-function mutations were manually introduced by substituting random cysteine binding residues (indicated with an asterisk) with non-polar alanines. The table depicts which metals MetaLATTE predicts for interaction, provided the input protein sequence specified.

## Discussion

Presently, regions characterized by intensive industrial operations, notably mining districts and densely populated urban areas, bear the brunt of heavy metal contamination, affecting nearly 40% of the Earth’s landmass [Tchounwou et al., 2012]. As a result, heavy metal-binding proteins may serve as bioremediation tools for metal contamination cleanup. As a first step toward this goal, in this work, we introduce a classifier that performs multi-label metal binding prediction for a given protein sequence. Our study highlights that classification outcomes depend heavily on label-specific thresholds. While we used EMA for threshold updates during training, further optimization will be needed. Nonetheless, our model produces improved representations for metal-binding proteins, especially those that bind to underrepresented metal ions, laying the groundwork for novel sequence design with controllable metal-binding features. Future work will include residue-specific predictors and visualization tools to identify key metal-binding residues, especially in proteins from understudied organisms. We aim to design low-complexity sequences rich in functional domains, mimicking natural metallothioneins. As these proteins tend to be structurally disordered [Calatayud et al., 2021] and are not amenable for structure prediction and design tools, such as AlphaFold3 [Abramson et al., 2024], we will continue to focus on sequence-based models for quality checks and novel designs. Overall, we envision that our model will enable the rapid screening of new metal-binding protein sequences, whether via metagenomic sequencing or *de novo* protein design, that bind to specified heavy metals, such as lead, cadmium, and mercury, for downstream bioremediation efforts. Finally, we hope that this study also raises awareness about the need to protect biodiversity and invest in basic biology, as the solutions to current environmental problems may lie within the understudied organisms yet to be discovered.

## Declarations

## Acknowledgements

We thank Mark III Systems and the Duke Computing Cluster for computing support. We further thank Willi Yu, Jing Guo, Xue Li, and Tianlai Chen for their insights related to the manuscript. The work was funded by a Garden Grant from the Homeworld Collective.

## Author Contributions

Y.Z. designed and implemented MetaLATTE architecture. Y.Z., P.H., and A.H. performed data preparation and model benchmarking. Y.Z. and P.C. wrote the manuscript. P.C. conceived, designed, directed, and supervised the study.

## Data and Materials Availability

All data needed to evaluate the conclusions are presented in the paper and tables. Model weights and code are located at https://huggingface.co/ChatterjeeLab/MetaLATTE.

## Supplementary Information

**Figure S1:**
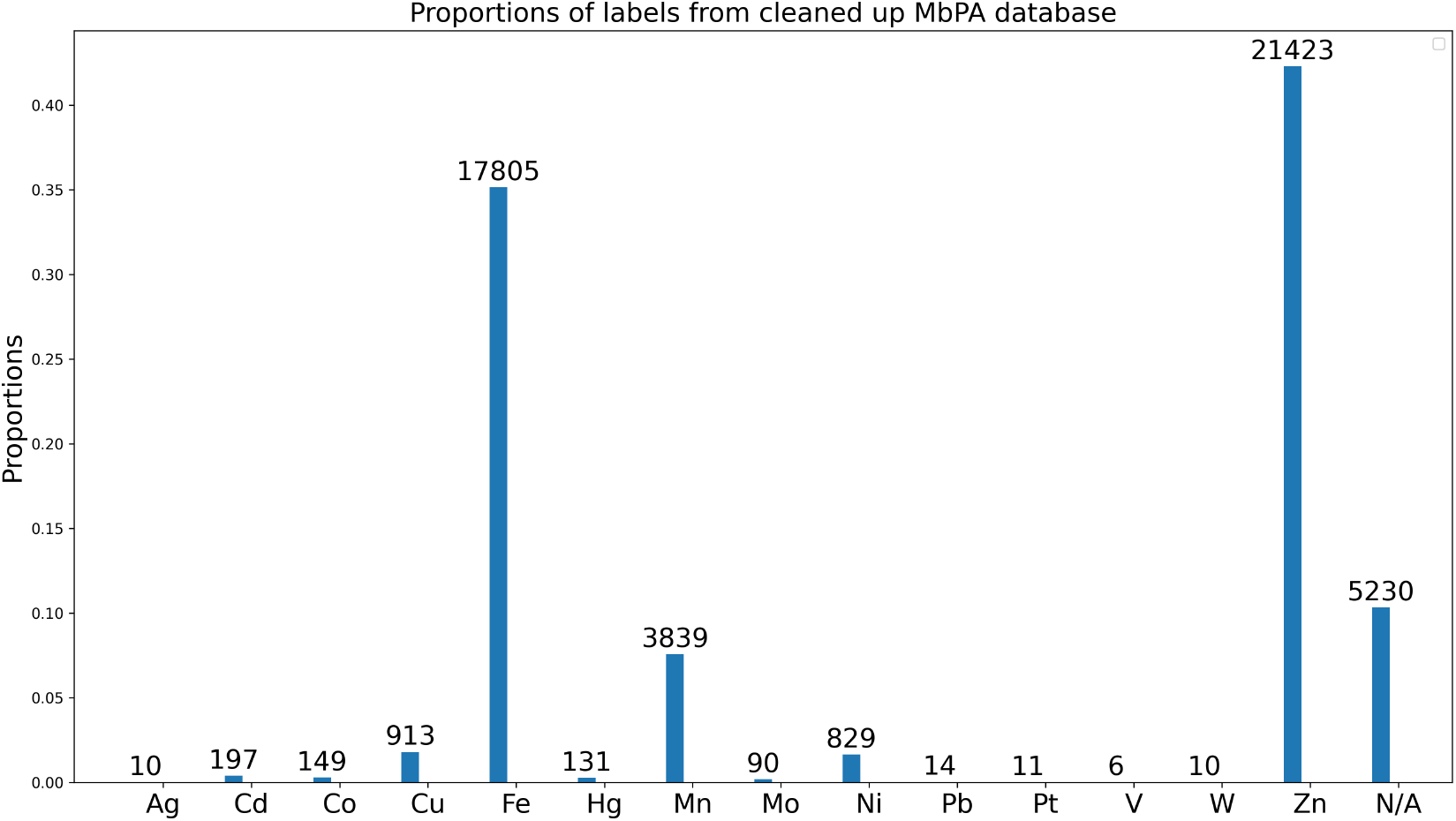
Label Proportions from MbPA database. The dataset was cleaned by filtering out non-transition metals and combining redundant sequence labels.

**Figure S2:**
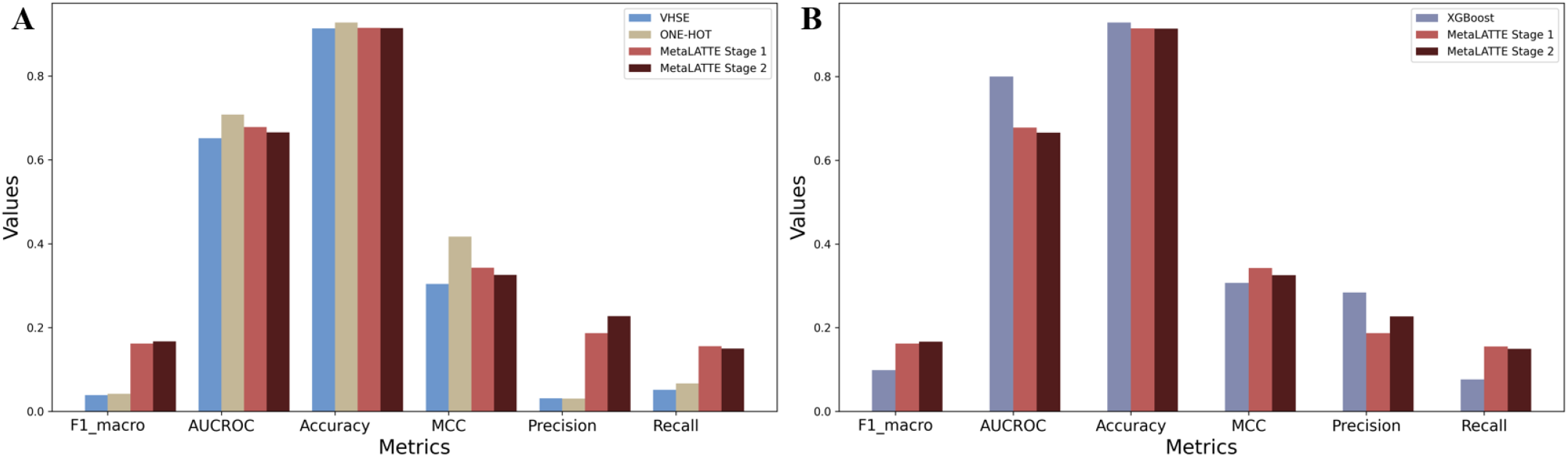
Multi-label classification metrics for the overall performance across all metal classes. (A) Benchmarking results for MetaLATTE against different protein embedding sources, including VHSE, one-hot, and ESM-2. (B) Benchmarking results for MetaLATTE among different models for processing ESM-2 embeddings, from left to right: XGBoost, MetaLATTE Stage 1, MetaLATTE Stage 2. The metrics evaluated on the x-axis are F1_macro, AUCROC, Accuracy, MCC, Precision, and Recall.

**Figure S3:**
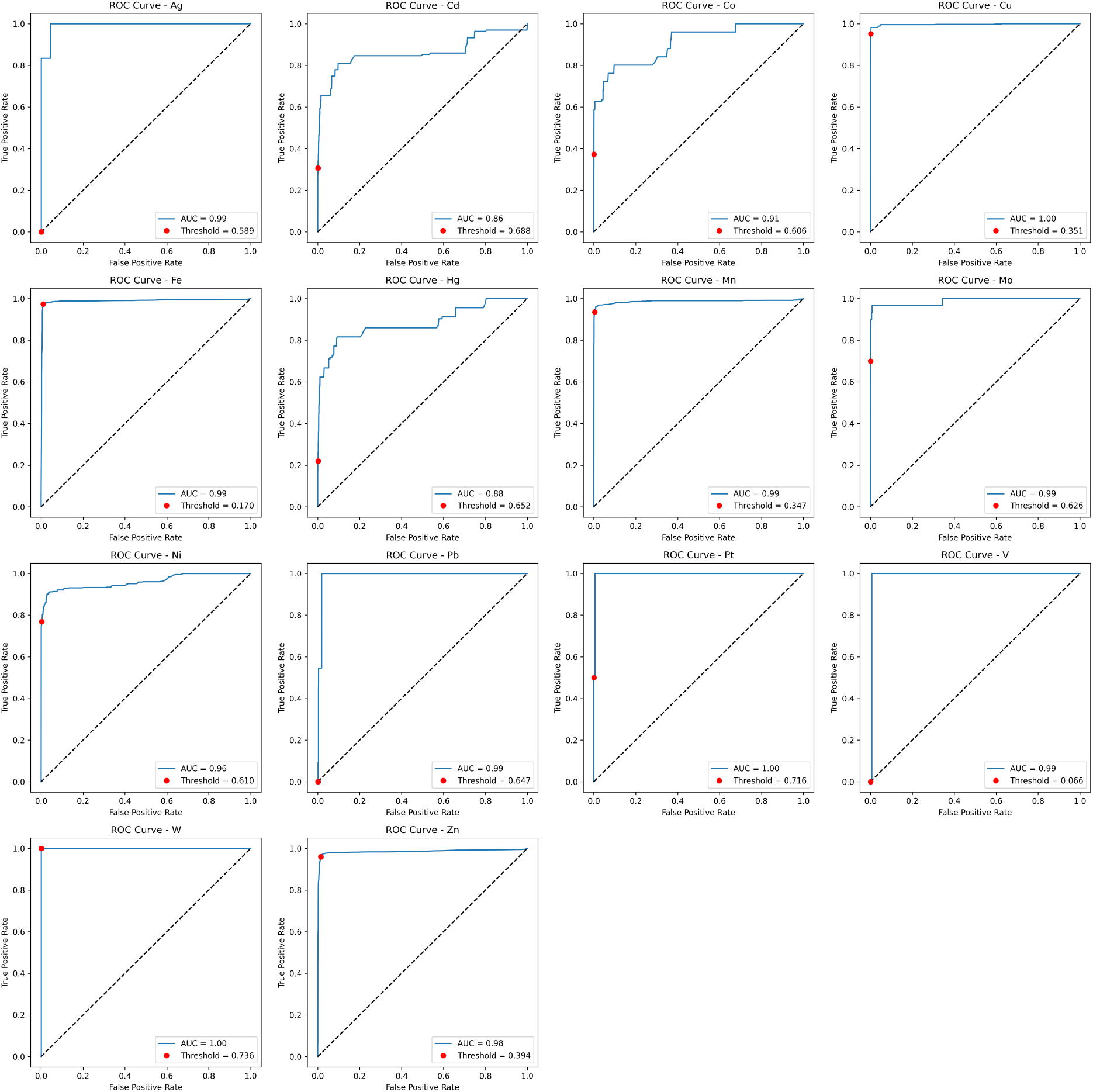
ROC curves for the MetaLATTE Stage 2 model with classification thresholds labeled as the red points per class. The metals classified are indicated in the plot titles.

**Figure S4:**
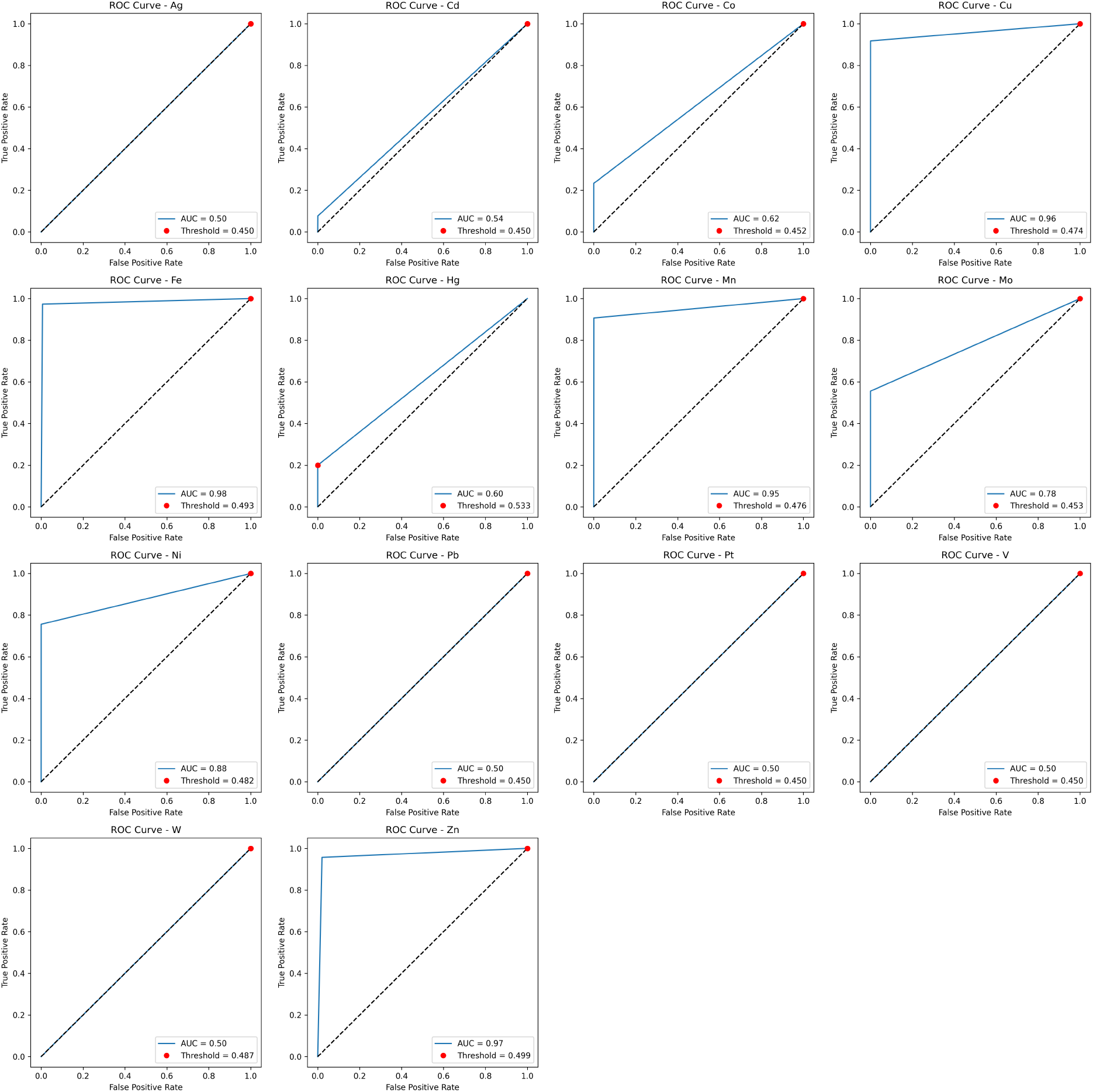
ROC curves for the optimized XGBoost model with classification thresholds labeled as the red points per class. The metals classified are indicated in the plot titles.

